# MICU1 modulates MCU ion selectivity and tolerance to manganese stress

**DOI:** 10.1101/371419

**Authors:** Jennifer Wettmarshausen, Valerie Goh, Utkarsh Tripathi, Anja Leimpek, Yiming Cheng, Alexandros A. Pittis, Toni Gabaldón, Dejana Mokranjac, Fabiana Perocchi

**Author notes:** Equal contribution (Jennifer Wettmarshausen, Valerie Goh). **Lead Contact**: Fabiana Perocchi.

## Abstract

The mitochondrial calcium uniporter is a highly selective ion channel composed of species-and tissue-specific structural and regulatory subunits. However, the contribution of each component to uniporter-mediated activity still remains unclear. Here, we employ an evolutionary and synthetic biology approach to investigate the functional inter-dependence between the pore-forming subunit MCU and the EF-hand protein MICU1. Using phylogenetic profiling and genetic complementation analyses, we show that MCU and MICU1 constitute the minimal eukaryotic unit of the uniporter, pointing towards a strong selective pressure behind their co-occurrence. Heterologous reconstitution of MCU-mediated and MICU1-gated mitochondrial calcium entry *in vivo* in yeast cells demonstrates that MICU1 *per se* is essential to protect yeast from MCU-dependent manganese cytotoxicity. Accordingly, MICU1 deletion significantly sensitizes human HEK-293 cells to manganese-induced stress. Our study identifies a critical role of MICU1 in the regulation of MCU ion selectivity, with potential implications for patients with MICU1 deficiency.

## INTRODUCTION

Intracellular calcium (Ca^2+^)-mediated signaling is an essential and universal mechanism to mediate adaptive physiological responses to a wide variety of environmental and endogenous stimuli, by regulating cell fate, differentiation, proliferation, metabolic flexibility, secretion, and gene expression (Berridge et al., 2000). Mitochondria from several organisms take an active role in the modulation of cytosolic calcium (cyt-Ca^2+^) dynamics. Due to their ability of rapidly and transiently accumulating Ca^2+^ and coupling it to ATP production (Denton, 2009; McCormack et al., 1990), mitochondria can shape, remodel, and decode intracellular Ca^2+^ signals and cover the energetic demand of signaling cells (Mammucari et al., 2018). Early studies on the properties of mitochondrial Ca^2+^ (mt-Ca^2+^) uptake in several vertebrate species have proposed that an electrophoretic uniporter mechanism is responsible for the transport of Ca^2+^ through the inner mitochondrial membrane, which makes use of the steep electrochemical gradient generated by the respiratory chain and is sensitive to general inhibitors of Ca^2+^-permeable ion channels (Carafoli and Lehninger, 1971; Deluca and Engstrom, 1961; Matlib et al., 1998; Vasington et al., 1972; Vasington and Murphy, 1962). This model was later confirmed by direct electrophysiological measurements of Ca^2+^ currents in single mammalian mitochondria (Fieni et al., 2012; Kirichok et al., 2004) and since then the physiological role of the mitochondrial calcium uniporter has been extensively investigated in many eukaryotic organisms (De Stefani et al., 2016).

Recently, a functional genomics approach combining comparative physiology analyses of mt-Ca^2+^ uptake, whole-genomes sequencing, and mitochondrial proteome mapping, has allowed the discovery of the first peripheral Ca^2+^-dependent regulator of mt-Ca^2+^ uptake (MICU1) (Perocchi et al., 2010) and the transmembrane pore-forming subunit of the mitochondrial calcium uniporter (MCU) (Baughman et al., 2011; De Stefani et al., 2011) in vertebrates and protozoa. Afterwards, several paralogs of MCU and MICU1, including the membrane-spanning dominant-negative regulator MCUB (Raffaello et al., 2013) and tissue-specific EF hand-containing subunits MICU2 and MICU3 (Patron et al., 2014; Patron et al., 2018; Plovanich et al., 2013), as well as the 10 kDa essential MCU interactor EMRE (Sancak et al., 2013), have been identified in Metazoa. Altogether, those proteins represent a toolkit of inhibitory and enhancing effectors of mt-Ca^2+^ uptake that fine-tune the activity of mammalian MCU in response to cell type and tissue-specific Ca^2+^ dynamics and energy requirements (De Stefani et al., 2016).

Overall, the complex molecular nature of the mammalian uniporter highlights the physiological relevance of achieving great plasticity and selectivity in Ca^2+^ uptake, by promoting Ca^2+^ entry over other divalent ions. Due to the presence of a very large driving force for cation influx, the uniporter must, at the same time, limit mt-Ca^2+^ accumulation when the cell is at rest, to prevent vicious Ca^2+^ cycling, but rapidly transmit a cyt-Ca^2+^ rise to the mitochondrial matrix during cell signaling. The highly selective permeability of the uniporter for Ca^2+^ is thought to derive from the high-affinity binding of the ion to the DXXE motif at the MCU pore (Arduino et al., 2017; Baughman et al., 2011; Cao et al., 2017; Chaudhuri et al., 2013; Oxenoid et al., 2016), whereas both gating and cooperative activation of the uniporter have been attributed to its interaction with hetero-oligomers of MICU1 and MICU2 or MICU3 (Csordas et al., 2013; Kamer et al., 2017; Mallilankaraman et al., 2012; Patron et al., 2014; Patron et al., 2018). However, the respective functional and mechanistic roles of those subunits in regulating uniporter activity have been so far investigated in mammalian systems where the interpretation of results is hampered by differences in the degree of gene silencing and tissue-specific protein composition (Murgia and Rizzuto, 2015; Vecellio Reane et al., 2016), stoichiometry and compensatory remodeling (Liu et al., 2016; Paillard et al., 2017) of the channel. Instead, the budding yeast *S. cerevisiae* represents an ideal testbed for dissecting the functional contribution of each exogenous component of the human uniporter, given that it completely lacks any detectable MCU homolog (Bick et al., 2012; Cheng and Perocchi, 2015) and endogenous mt-Ca^2+^ transport activity (Arduino et al., 2017; Carafoli and Lehninger, 1971; Kovacs-Bogdan et al., 2014; Yamamoto et al., 2016), while enabling the facile expression and targeting of human mitochondrial proteins. We and others have recently shown that the co-expression of human MCU and EMRE in yeast is sufficient to regulate the entry of a bolus of Ca^2+^ into the matrix of isolated mitochondria (Arduino et al., 2017; Kovacs-Bogdan et al., 2014; Yamamoto et al., 2016) with the same biophysical properties as the mammalian uniporter.

Here we employ yeast as a model system to investigate the functional relationship between human MCU and MICU1. Using a comparative genomics approach, we show that MCU and MICU1 constitute the minimal conserved unit of the uniporter throughout eukaryotic evolution, pointing towards a strong selective pressure behind their co-occurrence. We show that heterologous expression of human MCU, EMRE, and MICU1 are sufficient to reconstitute a Ca^2+^-regulated and uniporter-dependent uptake of Ca^2+^ from the cytosol to the mitochondria of yeast *in vivo* in response to environmental stimuli. We exploit this new experimental system to investigate MCU-MICU1 genetic interaction by screening for biological conditions whereby the co-occurrence of MCU and MICU1 would provide a selective fitness advantage. We find that MICU1 is *per se* essential to protect yeast cells from MCU-dependent iron (Fe^2+^) and manganese (Mn^2+^) toxicity, indicating an unanticipated regulatory function in the ion selectivity of the uniporter. We confirm the physiological relevance of our findings in human HEK-293 cells where the knock-out (KO) of MICU1 greatly sensitizes to Mn^2+^-induced cell death, which we rescue by re-introducing a wild-type (WT) or EF-hands mutant MICU1 as well as by exogenous Fe^2+^ supplementation. Altogether, our findings highlight a previously unknown regulatory role of MICU1 in MCU ion permeability, with potential implications for MICU1-related human disorders.

## RESULTS

### MCU and MICU1 form the minimal eukaryotic unit of the mitochondrial calcium uniporter

The molecular identification of the mitochondrial calcium uniporter complex, together with the availability of numerous sequenced genomes from diverse taxonomic groups and of genome annotation tools, offers the opportunity to exploit evolutionary genomics approaches to investigate the functional links between members of this channel. Correlated patterns of presence and absence of two or more proteins of the channel across a large number of genomes (co-occurrence) may indicate strong evolutionary constrains for their functional inter-dependence (de Juan et al., 2013). Here, we examined the phylogenetic profiles and predicted mitochondrial co-localization of the two founding members of the uniporter, MCU and MICU1 (Baughman et al., 2011; De Stefani et al., 2011; Perocchi et al., 2010), across 247 fully-sequenced eukaryotic species from multiple taxonomic levels at different evolutionary distances, in order to maximize the resolution of coupled evolutionary patterns (**Fig 1A**; see also https://itol.embl.de/tree/774755176425021526503446) (Cheng and Perocchi, 2015). Similarly to a previous evolutionary analysis across 138 species (Bick et al., 2012), we observed a widespread distribution of MCU homologs in all major eukaryotic groups, present in nearly all Metazoa and Plants, but only in some Protozoa (e.g., *Trypanosoma cruzi* and *Leishmania major*) and Fungi (e.g., *Neurospora crassa* and *Aspergillus fumigatus*). Specifically, homologs of MCU have been apparently lost in all Apicomplexa (e.g., *Plasmodium falciparum*), mitochondrial-devoid, single-cell eukaryotes (e.g., *Entamoeba histolytica*, *Giardia lambia*, *E. cuniculi*), and Saccharomycota (e.g., *S. cerevisiae, S.pombe, C. glabrata*), (see https://itol.embl.de/tree/774755176425021526503446).

**Figure 1.**
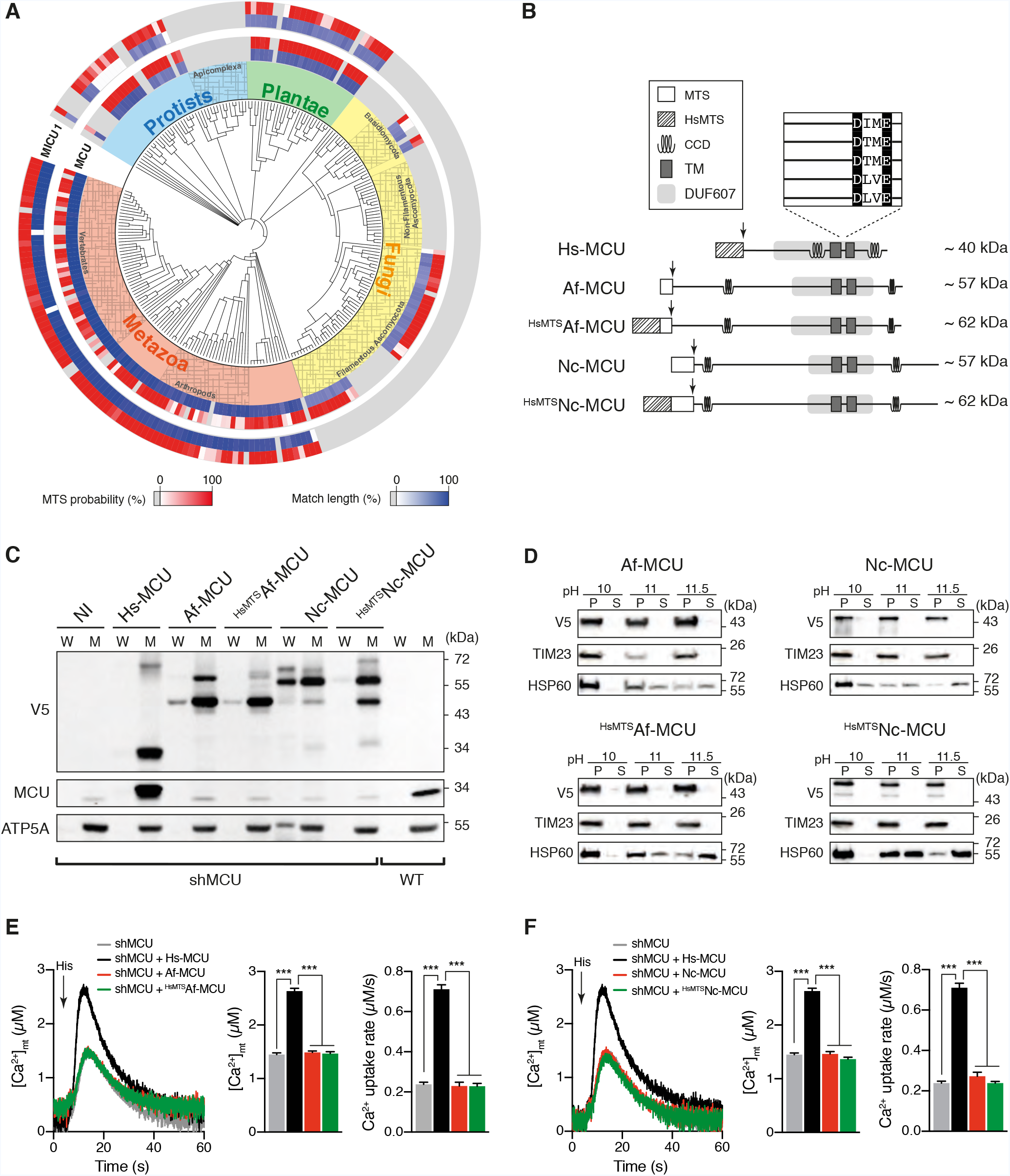
Phylogenetic profiling of MCU and MICU1. **(A)** Phylogenetic distribution of MCU and MICU1 homologs (blue, percentage of amino acids match length) across 247 eukaryotes. MTS, mitochondrial targeting sequence from MitoProt. **(B)** Schematic of ectopically expressed fungal MCU constructs and protein domains (CCD, coiled-coil domain; TM, transmembrane domain). DXXE motif and MTS cleavage site prediction (arrow) are also shown. **(C)** Analysis of whole cell (W) and mitochondrial (M) fractions isolated from MCU knocked-down (shMCU) HeLa cells stably expressing human (Hs-MCU), *N. crassa* (Nc-MCU), or *A. fumigatus* (Af-MCU) MCU fused with a C-terminal V5-tag and compared to wild-type (WT). ^HsMTS^Af-MCU and ^HsMTS^Nc-MCU refer to *A. fumigatus* and *N. crassa* MCU constructs fused at the N-terminal to the MTS of Hs-MCU (HsMTS). NI, not infected. **(D)** Analysis of mitochondrial soluble (S) and membrane pellet (P) fractions isolated from shMCU HeLa cells expressing Af-MCU and Nc-MCU with and without HsMTS. (**E**, **F**) Representative traces and quantification of mitochondrial Ca^2+^ kinetics in shMCU HeLa cells expressing *A. fumigatus* (E) or *N. crassa* (F) MCU constructs upon histamine (His) stimulation (100 μM final concentration). All data represent mean ± SEM; n=6-8; ***p < 0.0001, one-way ANOVA with Tukey’s Multiple Comparison Test.

Strikingly, the distribution of MICU1 homologs was largely overlapping with that of MCU (**Fig 1A**), pointing to a strong functional inter-dependence between the two proteins, which we now know to be part of the same complex. However, the existence of organisms within Basidiomycota and Ascomycota fungal clades with just MCU, such as *N. crassa* and *A. fumigatus* (Bick et al., 2012; Prole and Taylor, 2012) suggests that MICU1 can be dispensable for mt-Ca^2+^ homeostasis. Alternatively, it is possible that MCU-like genes found in Fungi do not represent functional homologs of the mammalian uniporter. The latter hypothesis is consistent with the observation that human and fungal MCU homologs show very low sequence similarity, which is mainly due to the presence of a generic DUF607 motif and two coiled coil domains (Finn et al., 2016). Furthermore, given the complete loss of EMRE in Fungi (Sancak et al., 2013), fungal MCU homologs should be self-sufficient to drive mt-Ca^2+^ uptake, just like the MCU from *Dictostillium discoideum* (Arduino et al., 2017; Kovacs-Bogdan et al., 2014). Instead, mitochondria from *N. crassa* and *A. fumigatus* were shown to lack Ca^2+^ uptake activities similar to the fast and high capacity MCU-dependent Ca^2+^ entry found in mammalian mitochondria (Carafoli and Lehninger, 1971; Gonçalves et al., 2015; Song et al., 2016).

To gain further insights into the functionality of fungal MCU homologs, we tested their ability to complement MCU loss-of-function in human cells. We expressed *A. fumigatus* (Af-MCU), *N. crassa* (Nc-MCU) or human MCU (Hs-MCU) with a C-terminal V5 tag in MCU knocked-down (shMCU) HeLa cells (**Fig 1B**). To ensure the targeting of fungal MCUs to human mitochondria, we also tested chimera proteins consisting of the Hs-MCU mitochondrial targeting sequence (HsMTS) fused to the full-length form of Nc-MCU (^HsMTS^Nc-MCU) and Af-MCU (^HsMTS^Af-MCU). We confirmed that all constructs were enriched in the mitochondria of shMCU HeLa cells (**Fig 1C**), similarly to a known mitochondria-localized protein (ATP5A) that was used as a control. Furthermore, all constructs showed proper insertion into the mitochondrial membrane from alkaline carbonate extraction analyses, compared to the integral inner membrane and soluble matrix targeted proteins TIM23 and HSP60, respectively (**Fig 1D**). Next, we quantified mt-Ca^2+^ uptake kinetics upon stimulation with histamine, a Ca^2+^ activating agonist, which we monitored with the luminescent Ca^2+^ sensor aequorin stably expressed in the mitochondrial matrix of HeLa cells (mt-AEQ). Although the expression of HsMCU in shMCU HeLa cells resulted in a significant rescue of mt-Ca^2+^ level and uptake rate, neither Af-MCU (**Fig 1E**) nor Nc-MCU (**Fig 1F**), with and without HsMTS, were able of rescuing the mt-Ca^2+^ uptake phenotype of shMCU HeLa cells.

Collectively, phylogenetic and genetics complementation analyses provide evidence for the lack of a functional human MCU homolog in Fungi (*A. fumigatus* and *N. crassa*), and point to another, albeit unknown, function of fungal DUF607-containing proteins. While the possibility of mt-Ca^2+^ uptake occurring in other filamentous fungi cannot be excluded, our findings strongly suggest that MCU and MICU1 form the minimal eukaryotic unit of the mitochondrial calcium uniporter.

### *In vivo* reconstitution of mitochondrial calcium uptake in yeast

We employed the yeast *S. cerevisiae* as a model system to gain insight into the evolutionary pressure behind the tight co-occurrence of MCU and MICU1 across eukaryotic organisms. Yeast is able to maintain its cyt-Ca^2+^ at sub-micromolar levels, even in the presence of external Ca^2+^ up to 100 mM, and can transiently increase it to activate calcineurin-dependent pro-survival, adaptive transcriptional responses to diverse environmental stresses (Cyert, 2003). We thus asked whether MCU-mediated mt-Ca^2+^ uptake could be reconstituted in yeast under physiological conditions that trigger a cyt-Ca^2+^ burst. To this goal, we tested the ability of Hs-MCU and Hs-EMRE to reconstitute mt-Ca^2+^ uptake in intact yeast cells in response to stimulus-induced cyt-Ca^2+^ dynamics (**Fig 2**). As a stimulus we used glucose-induced calcium (GIC) activation (**Fig 2A**), whereby the addition of a carbon source like glucose to hexoses-starved yeast cells activates a high affinity Ca^2+^ influx system (HACS) at the plasma membrane and allows the rapid entry of Ca^2+^ from the extracellular environment (Groppi et al., 2011).

**Figure 2.**
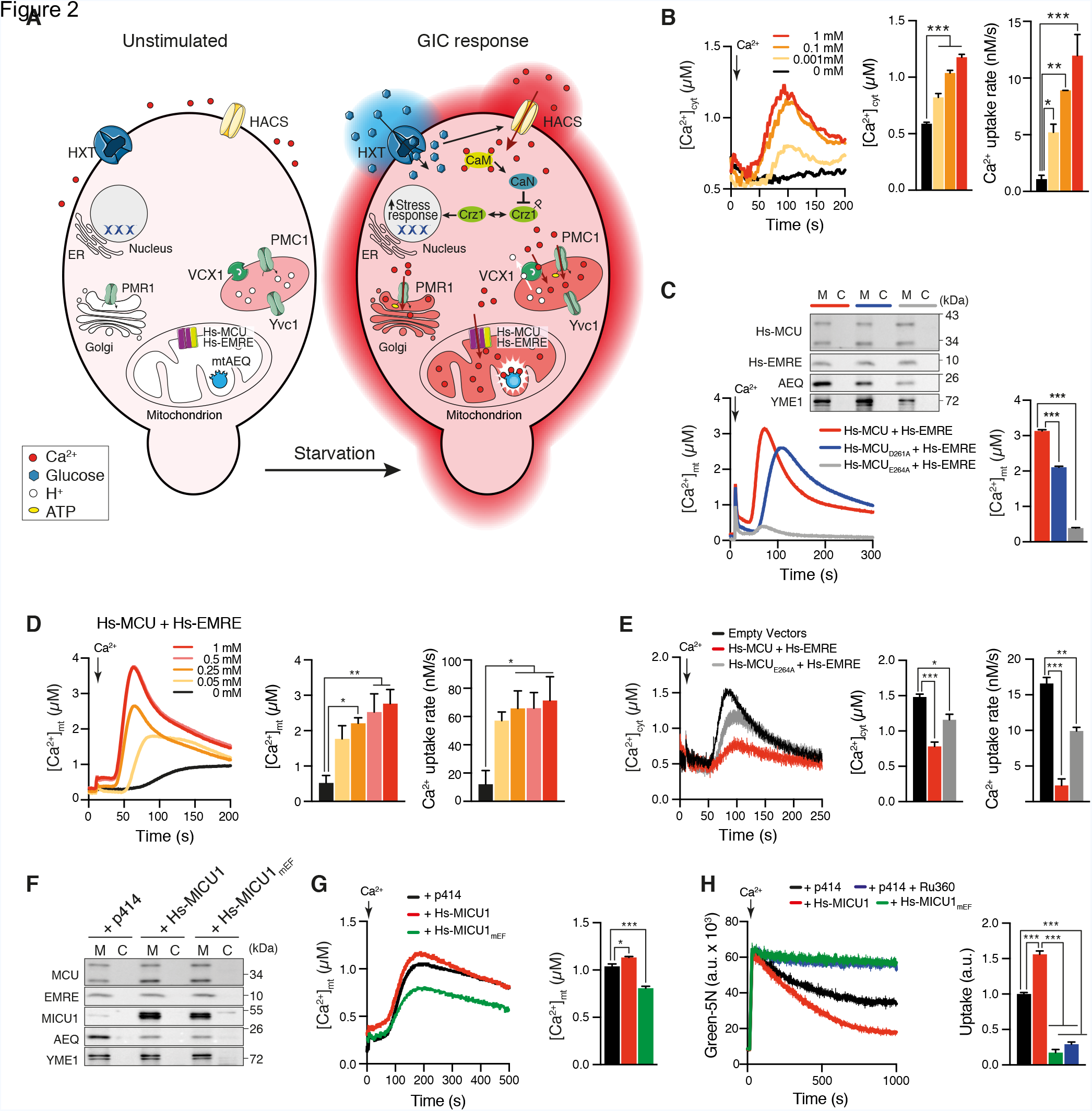
*In vivo* reconstitution of mitochondrial calcium uptake in yeast. **(A)** Schematic of the Ca^2+^ homeostasis system and glucose-induced calcium (GIC) signaling in *S. cerevisiae*. Hs-MCU, human MCU; Hs-EMRE, human EMRE; HXT, hexose transporter; HACS, high-affinity Ca^2+^ transport system; VCX1, vacuolar H^+^/Ca^2+^ exchanger; PMC1, vacuolar Ca^2+^-ATPase; PMR1, ER/golgi Ca^2+^-ATPase; mtAEQ, mitochondria-targeted aequorin; YVC1, TRPC-type Ca^2+^ channel; CaN, calcineurin; CaM, calmodulin; Crz1, calcineurin-dependent transcription factor; Crz1^P^, phosphorylated Crz1. **(B)** Dynamics and quantification of cytosolic Ca^2+^ kinetics in yeast cells expressing a cytosolic aequorin (cyt-AEQ) upon GIC stimulation in presence of different extracellular CaCl_2_ concentrations (n=3); *p < 0.05, **p <0.01, ***p<0.001, one-way ANOVA with Dunnett’s Multiple Comparison Test. **(C)** Dynamics and quantification of mitochondrial Ca^2+^ kinetics in yeast cells expressing mtAEQ, Hs-EMRE, and either wild type or mutated Hs-MCU upon GIC stimulation in presence of 1 mM CaCl_2_ (n=3); ***p < 0.0001, one-way ANOVA with Tukey’s multiple comparisons test. *Inset*, immunoblot analysis of cytosolic (C) and mitochondrial (M) fractions. **(D)** Dynamics and quantification of mitochondrial Ca^2+^ kinetics in yeast cells expressing mtAEQ, Hs-EMRE and Hs-MCU upon GIC stimulation in presence of different extracellular CaCl_2_ concentrations (n=3); *p < 0.05, **p <0.01, one-way ANOVA with Dunnett’s multiple comparisons test. **(E)** Dynamics and quantification of cytosolic Ca^2+^ kinetics in yeast cells expressing cyt-AEQ with either empty vectors (p425, p423) or Hs-EMRE and wild type or mutant Hs-MCU upon GIC stimulation in presence of different extracellular CaCl_2_ concentrations (n=3); *p < 0.05, **p < 0.01, ***p < 0.001, one-way ANOVA with Dunnett’s multiple comparisons test. **(F)** Immunoblot analysis of cytosolic (C) and mitochondrial (M) fractions isolated from yeast strains expressing mtAEQ, Hs-MCU, Hs-EMRE and either an empty vector (p414), human wild-type MICU1 (Hs-MICU1), or EF-hands mutant MICU1 (Hs-MICU1_mEF_). **(G)** Dynamics and quantification of mitochondrial Ca^2+^ kinetics in the yeast strains described in (F) upon GIC stimulation in presence of 1 mM CaCl_2_ (n=3); *p < 0.05, ***p < 0.001, one-way ANOVA with Dunnett’s multiple comparisons test. (**H**) Representative traces and quantification of extracellular Ca^2+^ clearance by mitochondria isolated from the yeast strains described in (F) upon addition of 100 μM final concentration of CaCl_2_ (n=3); ***p < 0.0001, one-way ANOVA with Tukey’s Multiple Comparison Test. 10 μM Ru360 (specific MCU inhibitor) was added as a positive control. All data represent mean ± SEM.

As a first step, we confirmed GIC-dependent activation of cyt-Ca^2+^ signaling by engineering a yeast strain that expresses a cytosol-localized aequorin (Nakajima-Shimada et al., 1991) and quantified cyt-Ca^2+^ kinetics in response to the re-addition of glucose to cells exponentially growing in a non-fermentative medium (**Fig 2B**). In agreement with previous results (Groppi et al., 2011), we observed that both the amplitude and uptake rate of cyt-Ca^2+^ responses were dependent on the external concentration of Ca^2+^. Next, we quantified GIC-induced mt-Ca^2+^ uptake using a yeast strain that we previously engineered to express mt-AEQ (Arduino et al., 2017). We showed that the heterologous co-expression of WT, full-length Hs-MCU and Hs-EMRE in yeast is essential to drive mt-Ca^2+^ uptake *in vivo* in response to glucose and in the presence of 1 mM external Ca^2+^ (**Fig 2C**). Accordingly, yeast strains expressing Hs-EMRE together with Hs-MCU mutants in highly conserved acidic residues within the DXXE motif (Hs-MCU_D261A_; Hs-MCU_E264A_) were either partially functional (Hs-MCU_D261A_) or almost completely unable (Hs-MCU_E264A_) to fully transfer GIC-induced cyt-Ca^2+^ signals into the mitochondrial matrix (**Fig 2C**). Moreover, MCU-mediated mt-Ca^2+^ uptake was responsive to a wide dynamic range of external Ca^2+^ concentrations (**Fig 2D**) and able to promptly buffer the rise of cyt-Ca^2+^ induced by glucose at 1 mM extracellular Ca^2+^ level (**Fig 2E**).

Afterwards, we tested whether the expression of human MICU1 in the yeast model system would be sufficient to reconstitute a channel that is regulated by Ca^2+^. We generated a yeast strain expressing Hs-MCU and Hs-EMRE together with either human WT (Hs-MICU1) or EF-hands mutant (Hs-MICU1_mEF_) MICU1 and showed that those were properly localized to mitochondria (**Fig 2F**). Notably, the presence of Hs-MICU1 significantly increased MCU-dependent mt-Ca^2+^ uptake in response to GIC signaling compared to the strain only expressing Hs-MCU and Hs-EMRE, whereas Hs-MICU1_mEF_ exerted an inhibitory effect (**Fig 2G**), as measured by mtAEQ-based light kinetics. To further confirm these results, we also quantified mt-Ca^2+^ buffering capacity in isolated yeast mitochondria using calcium green-5N as a Ca^2+^-sensitive fluorescent probe. Consistently with previous results (Yamamoto et al., 2016), measurements of Ca^2+^ clearance showed that mitochondria from a yeast strain expressing Hs-MCU, Hs-EMRE and Hs-MICU1 were more efficient in buffering a bolus of high Ca^2+^ (100 μM) from the extramitochondrial space when compared to the same strain without Hs-MICU1 (**Fig 2H**). As expected, the expression of Hs-MICU1_mEF_ dramatically reduced mt-Ca^2+^ uptake, similar to the effect of the MCU inhibitor Ru-360. Altogether, these results are consistent with findings in mammalian cells, where functional MICU1 Ca^2+^-sensing domains are necessary to activate MCU at high cyt-Ca^2+^ levels (Csordas et al., 2013; Kamer et al., 2017; Mallilankaraman et al., 2012; Patron et al., 2014; Patron et al., 2018).

Finally, our results demonstrate that human MCU and EMRE are sufficient to reconstitute mt-Ca^2+^ uptake *in vivo* in response to physiological stimuli that trigger cyt-Ca^2+^ signaling. They also show that the presence of human MICU1 in yeast can regulate mt-Ca^2+^ uptake and it is *per se* sufficient to confer a positive synergistic effect on MCU-mediated mt-Ca^2+^ kinetics.

### MCU impairs yeast tolerance to iron and manganese stress

Having reconstituted mammalian-like mt-Ca^2+^ signaling *in vivo* in yeast, we sought to exploit this result to search for biological conditions where the presence of MICU1 would confer a fitness advantage. We hypothesized that a strain expressing only Hs-MCU and Hs-EMRE, which is able to buffer cyt-Ca^2+^ elevations (**Fig 2E**), should be more sensitive to environmental stresses, given that fungal cells need cyt-Ca^2+^ transients to activate pro-survival responses (Cyert, 2003). To test our hypothesis, we compared the fitness of a WT strain to that of yeast strains expressing Hs-EMRE together with either functional (Hs-MCU) or inactive MCU (Hs-MCU_E264A_) under an array of different environmental conditions, including heat shock, fungicide treatment, high salt and heavy metals stresses (Bultynck et al., 2006; Cyert, 2003; Edlind et al., 2002; Farcasanu et al., 1995; Gupta et al., 2003; Heuck et al., 2010; Nakamura et al., 1993; Pagani et al., 2007; Peiter et al., 2005; Serrano et al., 2004), by employing growth rate as a proxy for cell survival and proliferation (**Fig 3**). To ensure the reliance of cell proliferation on fully developed and functional mitochondria, all strains were tested during exponential growth in a non-fermentable medium, with lactate as a carbon source. Overall, we observed comparable doubling times among the three different strains during normal growth at 30 °C in lactate medium, which was over two-fold higher upon heat shock (37 °C) (**Fig 3A**). Likewise, treatment with increasing doses of two antifungal drugs, miconazole and amiodarone, either decreased the growth rate of the yeast cultures by over two-fold (miconazole, 100 ng/ml) (**Fig 3B**) or resulted in a complete cessation of growth (amiodarone, 20 μM) (**Fig 3C**), independent of the genetic background. All three strains also showed similar sensitivities to high-salt stress when grown under increasing concentration of NaCl (**Fig 3D**).

**Figure 3.**
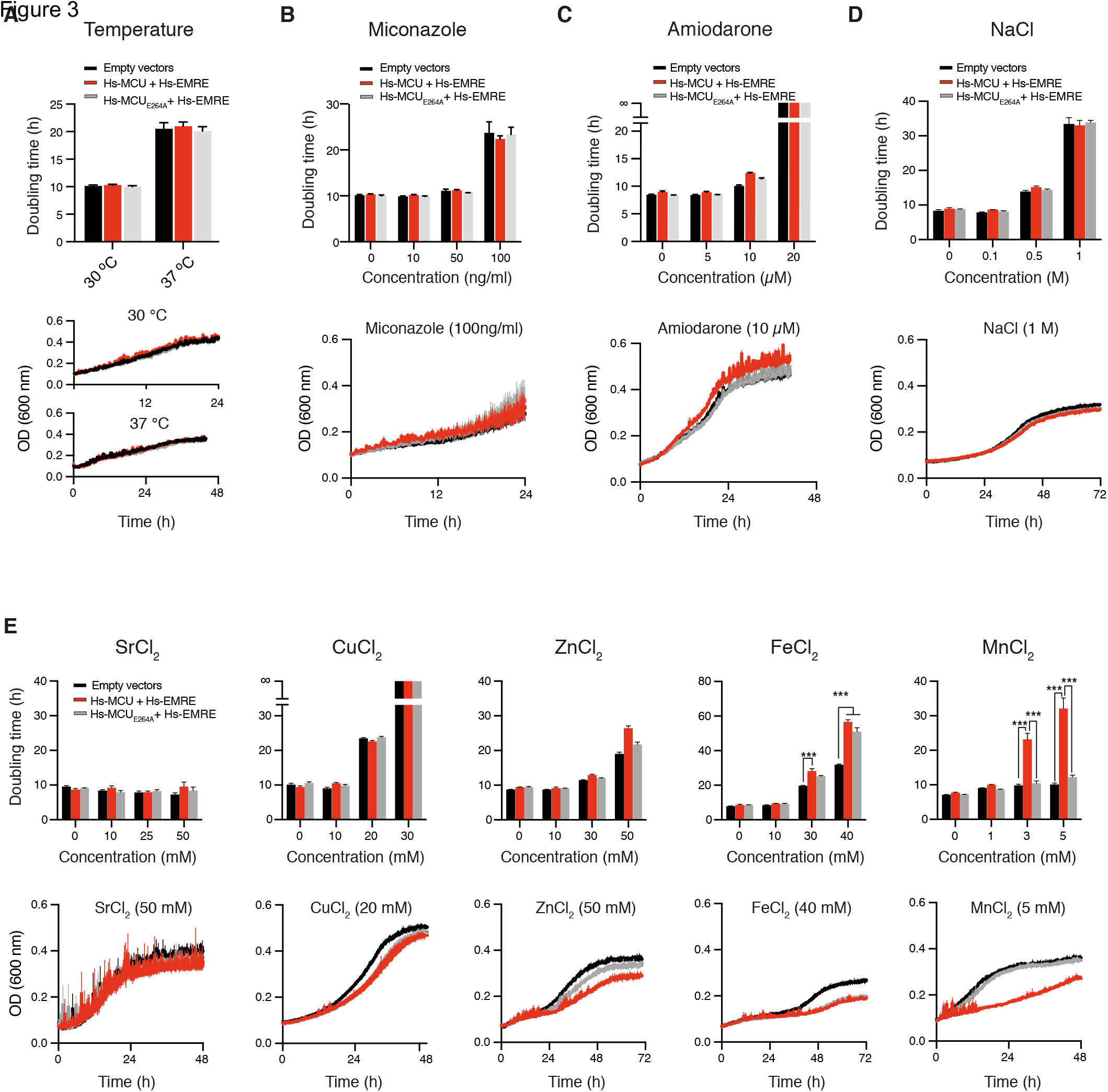
MCU impairs yeast tolerance to iron and manganese stress. (**A-E**) Quantification of growth rate and average growth curve of yeast strains expressing empty vectors (p423 and p425) or Hs-EMRE with wild-type or mutated Hs-MCU at 30°C and 37°C (A), and at increasing concentrations of miconazole (B), amiodarone (C), NaCl (D), or heavy metals (E). SrCl_2_, strontium chloride; CuCl_2_, copper chloride; ZnCl_2_, zinc chloride; FeCl_2_, iron chloride; MnCl_2_, manganese chloride. Data represent mean ± SEM; n=4; ***p < 0.0001, one-way ANOVA with Tukey’s Multiple Comparison Test.

Instead, we observed notable different responses among strains to heavy metals-induced stress (Sr^2+^, Cu^2+^, Zn^2+^, Fe^2+^, Mn^2+^) (**Fig 3E**). Although minimal levels of those cations are important for normal metabolism, high concentrations can be toxic (Farcasanu et al., 1995; Heuck et al., 2010; Pagani et al., 2007; Peiter et al., 2005; Serrano et al., 2004). Accordingly, at extracellular concentrations of CuCl_2_ and ZnCl_2_ above 10 and 30 mM, respectively, the doubling time of yeast cultures was over two-fold higher, with the exception of Sr^2+^, for which concentrations above 50 mM could not be tested due to precipitation in the liquid medium. However, while these three strains did not show any significant difference in their tolerance to those cations, we observed a greater hypersensitivity of the functional MCU-reconstituted strain to both Fe^2+^ and Mn^2+^ toxicity, which manifested with a drastic reduction in cell proliferation at concentrations above 10 and 1 mM, respectively (**Fig 3E**). Strikingly, the MCU_E264A_–expressing strain showed the same tolerance as the WT strain to Mn^2+^, whereas tolerance to Fe^2+^ was independent of a functional DXXE motif.

Altogether, these results confute our initial hypothesis that the reconstitution of MCU-dependent mt-Ca^2+^ buffering in yeast would increase yeast susceptibility to environmental stresses by antagonizing Ca^2+^-dependent calcineurin signal transduction cascades, given that the latter is a common link in all tested stress conditions. Instead, our results indicate that the differences in growth rate observed between WT and MCU-reconstituted yeast strains at high extracellular Fe^2+^ and Mn^2+^ levels could be due to the known permeability of MCU to those cations (Ernster, 1978; Hillered et al., 1983; Hughes and Exton, 1983; Medvedeva and Weiss, 2014; Mela and Chance, 1968; Romslo and Flatmark, 1973; Saris, 2012; Saris and Kaija, 1994; Vainio et al., 1970; Vinogradov and Scarpa, 1973), whose accumulation in mitochondria is known to trigger cell death (Smith et al., 2017; Sripetchwandee et al., 2014). Furthermore, whereas our findings corroborate previous evidence of Mn^2+^ binding and conductance by MCU (Cao et al., 2017), they suggest that the DXXE motif confers selectivity to some but not all cations.

### MICU1 protects yeast and human cells from MCU-dependent manganese toxicity

Given the dramatic reduction of yeast tolerance to Mn^2+^ stress in the presence of Hs-MCU, we tested whether the reconstitution of a Hs-MICU1 regulated uniporter could rescue this phenotype (**Fig 4**). Strikingly, the expression of either WT or EF-hands mutant MICU1 significantly rescued the hypersensitivity of the Hs-MCU and Hs-EMRE reconstituted strain towards Fe^2+^ (**Fig 4A**) and Mn^2+^ (**Fig 4B**) stress. These results indicate that the interaction of Hs-MICU1 with Hs-MCU and Hs-EMRE, but not its EF-hands, is sufficient to prevent both Fe^2+^ and Mn^2+^ accumulation into the mitochondrial matrix, perhaps by keeping the channel in a close conformation. We then sought to determine whether our findings of a MCU-mediated toxicity and MICU1-dependent protection towards Fe^2+^ and Mn^2+^ stress observed in yeast cells would be recapitulated in mammalian cells. To this goal, we compared the viability of WT and MICU1 knockout (MICU1-KO) HEK-293 cells upon treatment with increasing concentrations of either FeCl_2_ (**Fig 4C**) or MnCl_2_ (**Fig 4D**) for 48 hours. Contrary to yeast cells, the absence of MICU1 did not sensitise cells to Fe^2+^ stress, as there was no difference in the viability of WT and MICU1-KO cells. Instead, we found a significant increase of cell death in MICU1-KO cells treated with concentrations of MnCl_2_ that are not lethal to WT cells. As observed in yeast, the protective role of MICU1 towards Mn^2+^ toxicity was not dependent on having functional Ca^2+^-sensing domains, as the stable re-introduction of both WT and EF-hands mutant MICU1 in MICU1-KO cells significantly rescued cell viability at 25 μM Mn^2+^ (**Fig 4E**).

**Figure 4.**
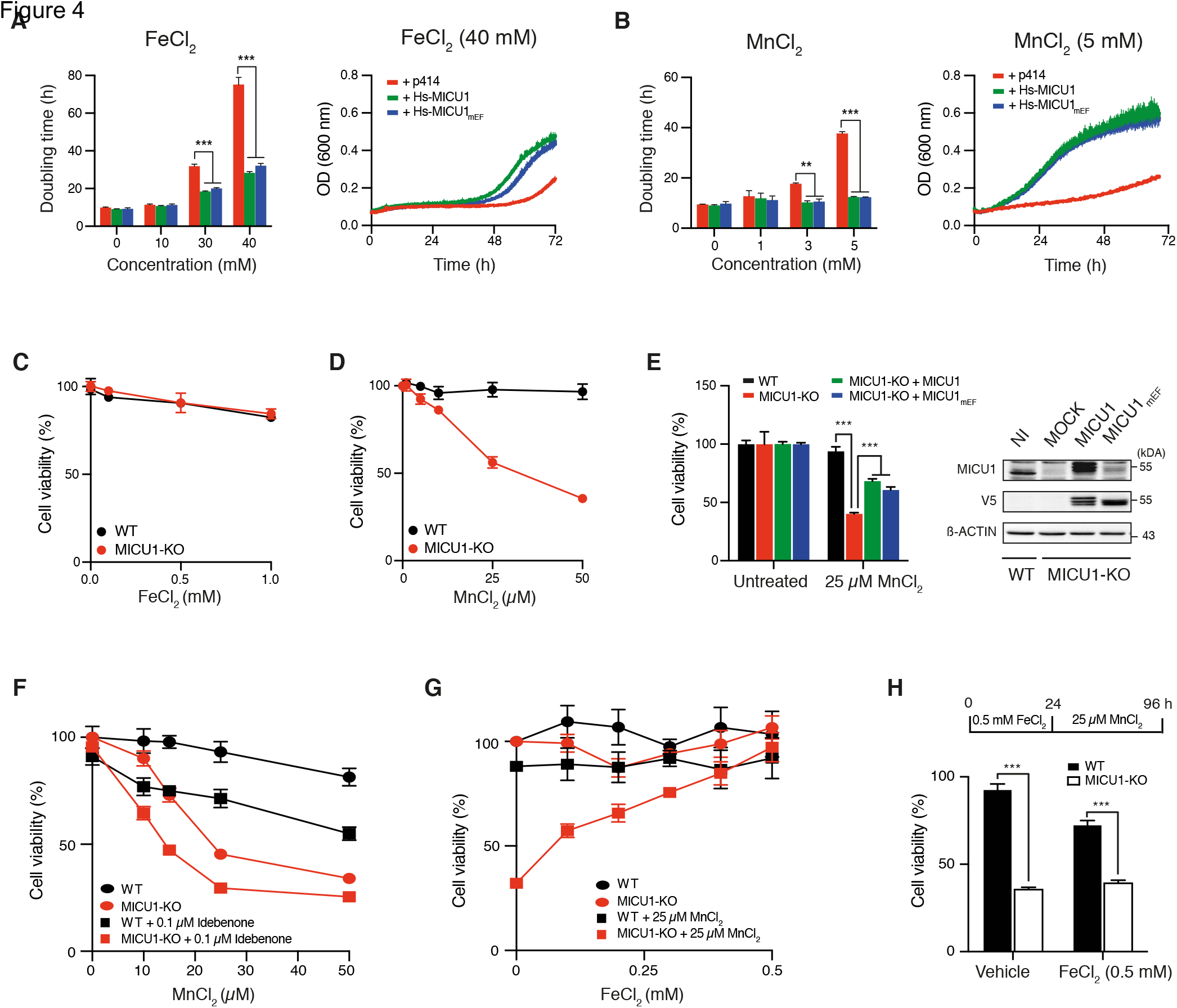
MICU1 protects yeast and human cells from manganese-induced cell death. (**A**, **B**) Quantification of growth rate and average growth curve of yeast strains expressing Hs-MCU, Hs-EMRE and either empty vector (p414), Hs-MICU1 or Hs-MICU1_mEF_ and treated with FeCl_2_ (A) or MnCl_2_ (B); n=4; ***p < 0.0001, one-way ANOVA with Tukey’s Multiple Comparison Test. (**C**, **D**) Cell viability assays of wild-type (WT) and MICU1 knockout (MICU1-KO) HEK-293 cells treated for 48 hours with FeCl_2_ (C) or MnCl_2_ (D); n=4. (**E**) Cell viability assays of uninfected WT (NI) and MICU1-KO cells that were either MOCK-treated or rescued by lentiviral expression of Hs-MICU1 or Hs-MICU1mEF upon MnCl_2_ treatment for 48 hours; n=4; ***p < 0.001, one-way ANOVA with Tukey’s Multiple Comparison Test. *Right*, immunoblot analysis of whole cell lysates from all cell lines. (**F**, **G**) Cell viability assays of WT and MICU1-KO HEK-293 cells treated for 48 hours with MnCl_2_ in the presence or absence of idebenone (n=4) or FeCl_2_ (n=3). (**H**) Cell viability assays of WT and MICU1-KO HEK-293 cells pre-treated with either FeCl_2_ or vehicle (water) for 24 hours followed by 48 hours of incubation with MnCl_2_ (n=3). All data represent mean ± SEM and are reported as the percentage of viable cells in untreated samples.

The above observations suggest that patients with MICU1 loss-of-function mutations (Lewis-Smith et al., 2016; Logan et al., 2014; Musa et al., 2018) might be more susceptible to accumulate Mn^2+^ in mitochondria, with potentially toxic effects (O’Neal and Zheng, 2015). We thus looked for strategies to ameliorate Mn^2+^ toxicity in MICU1-KO cells. Although the mechanism of mitochondrial Mn^2+^ toxicity is not entirely understood, it is believed that the major effect is oxidative stress, resulting in the induction of cell death (Smith et al., 2017). However, we could not detect a significant improvement in the viability of MICU1-KO cells exposed to high Mn^2+^ in the presence of 0.1 μM idebenone (**Fig 4F**), a clinically approved antioxidant drug used in mitochondrial OXPHOS disorders (Holzerova et al., 2016). As an alternative strategy, we co-treated MICU1-KO cells with Mn^2+^ in presence of 0.5 mM extracellular FeCl_2_ (**Fig 4G**), as both Fe^2+^ and Mn^2+^ compete for entry into cells via the divalent metal transporter 1 (DNM1) (Gunshin et al., 1997) and Fe^2+^ supplementation has been already shown to reduce Mn^2+^ overload and toxicity (O’Neal and Zheng, 2015; Tai et al., 2016). Consistently, we observed that the tolerance of MICU1-KO cells to Mn_2_ stress was completely restored, whereas pre-treatment with the same concentration of FeCl_2_ for 24 h was ineffective (**Fig 4H**). The latter confirms that the protective effect of Fe^2+^ supplementation results from reducing Mn^2+^ entry into the cell (O’Neal and Zheng, 2015; Tai et al., 2016).

In summary, these results establish a new functional role of MICU1 in conferring ion selectivity to the uniporter channel, which is independent of the EF-hands, but essential for preventing Mn^2+^ toxicity, with possible implications for patients with MICU1 deficiency.

## DISCUSSION

Our phylogenetic analyses, as well as a previous comparative genomics study (Bick et al., 2012), have highlighted a widespread co-occurrence of MCU and MICU1 across Metazoa, Plants and Protozoa, with the exception of Fungi. The presence of MCU homologs in several Ascomycota and Basidiomycota fungal clades devoid of a detectable MICU1, together with the absence of EF-hands containing proteins co-evolving with fungal MCUs (Cheng and Perocchi, 2015) and the existence of a MCU homolog that is sufficient to drive Ca^2+^ uptake in mitochondria of *D. discoideum* (Kovacs-Bogdan et al., 2014), led to the hypothesis that MCU could exist independently of a Ca^2+^-sensing regulator. However, this hypothesis stems from the assumption that fungal MCU homologs per se are able to mediate mt-Ca^2+^ uptake. Our results confute this hypothesis by demonstrating that fungal MCUs are not functional homologs of human MCU, given that they cannot complement the loss of uniporter-dependent Ca^2+^ uptake in mitochondria of MCU knockdown human cells (**Fig 1**). In light of these findings and previous evidence for the lack of a mammalian-like mt-Ca^2+^ uptake in filamentous fungi such as *N. crassa* and *A. fumigatus* (Carafoli and Lehninger, 1971; Gonçalves et al., 2015; Song et al., 2016), we propose that MCU and MICU1 constitute the minimal functional unit of the eukaryotic uniporter, and speculate that a strong selective pressure must have ensured their functional interdependence.

To investigate the direct contribution of MICU1 on MCU function, we employed the yeast *S. cerevisiae* as a model system, which lacks homologs of all mammalian uniporter components (Bick et al., 2012; Cheng and Perocchi, 2015), as well as endogenous mt-Ca^2+^ uptake activity (Arduino et al., 2017; Carafoli and Lehninger, 1971; Kovacs-Bogdan et al., 2014; Yamamoto et al., 2016). Previous results, including ours, have shown that Hs-MCU and Hs-EMRE are sufficient to drive Ca^2+^ uptake *in vitro* into the matrix of isolated mitochondria (Arduino et al., 2017; Kovacs-Bogdan et al., 2014; Yamamoto et al., 2016). While those analyses have corroborated the role of MCU as the *bona fide* pore-forming subunit of the uniporter, they did not assess its relevance for the efficient transfer of physiological Ca^2+^ transients from the cytosol to the mitochondrial matrix of yeast *in vivo* during signaling. We demonstrate that the heterologous expression of human MCU and EMRE can reconstitute mt-Ca^2+^ entry *in vivo* in yeast in response to a physiological rise in cyt-Ca^2+^ (**Fig 2**) and, similar to mammalian cells, the expression of human MICU1 in MCU-reconstituted yeast cells exerts a synergistic effect on mt-Ca^2+^ uptake kinetics, which is dependent on its Ca^2+^-sensing domains. Next, we searched for biological conditions whereby a positive MCU-MICU1 genetic interaction would provide a selective fitness advantage over a yeast strain reconstituted with MCU without its regulator. To this goal, we investigated the susceptibility of the MCU-expressing yeast strain to an array of stress conditions that require cyt-Ca^2+^ signaling for survival, including heat shock, salt, heavy metals-and fungicides-induced stresses (**Fig 3**). Surprisingly, we found that reconstitution of mtCa^2+^ uptake in yeast did not lead to a fitness disadvantage under standard non-fermentable growth conditions, but it was synthetically lethal with growth in the presence of high Mn^2+^ and Fe^2+^ concentrations in the extracellular medium.

The increased susceptibility of MCU-reconstituted yeast cells to Mn^2+^ and Fe^2+^ stresses is likely due to the permeation of those ions across the uniporter channel, which has been extensively documented (Ernster, 1978; Hillered et al., 1983; Hughes and Exton, 1983; Mela and Chance, 1968; Romslo and Flatmark, 1973; Saris, 2012; Saris and Kaija, 1994; Sripetchwandee et al., 2014; Vainio et al., 1970; Vinogradov and Scarpa, 1973), leading to their accumulation in mitochondria and the consequent toxicity (Smith et al., 2017; Sripetchwandee et al., 2014). This is consistent with the observation that the carboxylate rings formed by the D and E amino acids of the DXXE motif can bind Mn^2+^ in a cooperative and specific manner even in the presence of Ca^2+^ (Cao et al., 2017). Although, the direct interaction of other cations, such as Fe^2+^, with the selectivity filter of MCU has not yet been explored, our results indicate that at least the E of the DXXE motif is dispensable for MCU-mediated Fe^2+^ entry. Instead, we found that the presence of MICU1 in the MCU-reconstituted strain is sufficient to protect yeast cells against uniporter-dependent Mn^2+^ and Fe^2+^-induced toxicity (**Fig 4**). These findings were recapitulated in mammalian cells, where the knockout of MICU1 hypersensitized HEK-293 cells to Mn^2+^-dependent cell death. Here, Fe^2+^ treatment did not affect the cell viability of either MICU1-KO or WT cells, even at high non-physiological concentrations, suggesting that the absence of MICU1 is not sufficient to alter Fe^2+^ homeostasis in mammalian cells, where additional regulatory mechanisms might be in place to prevent cellular Fe^2+^ overload (Philpott, 2012). Notably, we found that functional MICU1 EF-hands were dispensable to confer protection against Mn^2+^ cytotoxicity, in contrast to their key role in enhancing Ca^2+^ uptake in response to a cyt-Ca^2+^ concentration above threshold (Csordas et al., 2013; Kamer et al., 2017; Mallilankaraman et al., 2012; Patron et al., 2014; Patron et al., 2018). Indeed, the Mn^2+^-induced toxicity observed in both MICU1-KO HEK-293 cells and MCU-reconstituted yeast strain was ameliorated by a stable expression of either WT or EF-hands mutant MICU1. This suggests that either Mn^2+^ does not target MICU1 EF-hands or its binding does not trigger the same conformational change that is elicited by Ca^2+^. The latter is consistent with previous studies demonstrating that Mn^2+^ can bind EF-hands without eliciting any protein conformational change, confirming their role also for ion selectivity, in addition to ion channel gating (Senguen and Grabarek, 2012; Shirran and Barran, 2009). Altogether, those results demonstrate that MICU1 is an important modulator of the ion selectivity of the mitochondrial calcium uniporter.

Several patients with loss-of-function mutations in MICU1 have been identified over the last few years, displaying a mixture of neurological and myopathic defects (Lewis-Smith et al., 2016; Logan et al., 2014; Musa et al., 2018), which were also recapitulated in MICU1-KO mice (Antony et al., 2016; Liu et al., 2016). Altogether, the disease phenotypes were attributed to a high basal mitochondrial Ca^2+^ level, possibly due to the loss of MICU1-dependent gatekeeping of the uniporter. Our results would now suggest that the increase in basal mt-Ca^2+^ level upon MICU1 loss-of-function could also result from the loss of MICU1-mediated protection towards Mn^2+^ entry into mitochondria. The latter would have a two-fold effect on mt-Ca^2+^ level and cell viability. First, Mn^2+^ accumulation into mitochondria is known to inhibit both the Na^+^-dependent and Na^+^-independent mitochondrial Ca^2+^ efflux routes (Gavin et al., 1990). Second, Mn^2+^ can block the uptake of Ca^2+^, Sr^2+^ and Ba^2+^, whereas pretreatment of mitochondria with Ca^2+^ increases the uptake of Mn^2+^ (Ernster, 1978; Hillered et al., 1983; Hughes and Exton, 1983; Mela and Chance, 1968; Vainio et al., 1970; Vinogradov and Scarpa, 1973). Last, while low doses of Mn^2+^ are necessary for normal cell and mitochondrial physiology, their accumulation is neurotoxic (Smith et al., 2017). Humans are often exposed to environmental sources of Mn^2+^, for example by consuming Mn^2+^-rich food such as grain products and vegetables, or by exposure to Mn^2+^ aerosols and dusts in mines, smelters, or simply to air pollution from the combustion of gasoline containing methylcyclopentadienyl manganese tricarbonyl (O’Neal and Zheng, 2015). Currently, adjuvant treatments include a dietary restriction of daily Mn^2+^ intake as well as Fe^2+^ supplementation (O’Neal and Zheng, 2015). Indeed, we also observed that co-treatment of Mn^2+^-stressed MICU1-KO cells with Fe^2+^ was able to prevent Mn^2+^ cytotoxicity in a dose-dependent manner, whereas pre-treatment with Fe^2+^, even at high doses, was without any effects. It can be speculated that human mutations in MICU1, that are loss of function for the protein activity, also affect MCU permeability to the heavy metals, and to Mn^2+^ in particular. Whether this altered ion permeability contributes to the phenotype of the patients remains to be demonstrated.

In summary, our study demonstrates the power of combining comparative genomics analyses with the use of yeast as a model system for dissecting the functional and mechanistic role of each component of the mammalian uniporter. Moreover, our reconstitution of a MICU1-regulated uniporter in yeast offers an incomparable advantage over similar investigation of MICU1 and MCU inter-dependence in mammalian cells, where MICU1 knockout or knockdown has also confounding effects on the expression of other uniporter subunits, such as MICU2 and MICU3 (Patron et al., 2014; Patron et al., 2018; Plovanich et al., 2013). Altogether, our results indicate that MICU1 is not only required for coupling mt-Ca^2+^ entry to changes in cyt-Ca^2+^ concentrations, but it also regulates the uniporter selective gating to Ca^2+^ ions, with important implications for patients with MICU1 deficiency.

## EXPERIMENTAL PROCEDURES

Additional details and resources used in this work can be found in Supplemental Experimental Procedures.

### Phylogenetic Profiling

The phylogenetic profiles of MCU and MICU1 across 247 eukaryotic species were generated using ProtPhylo (www.protphylo.org) (Cheng and Perocchi, 2015). Protein sequences of *Homo sapiens* MCU (Hs-MCU, NP_612366.1) *Neurospora crassa* MCU (Nc-MCU, XP_959658.1), and *Aspergillus fumigatus* MCU (Af-MCU, XP_751795.1) were analyzed to predict protein domains.

### Cell Lines

All mammalian cells were grown in high-glucose Dulbecco’s modified Eagle’s medium (DMEM) supplemented with 10% FBS, 100 μg/ml geneticin (mt-AEQ HeLa), 2 μg/mL puromycin (shMCU HeLa), 10 μg/mL blasticidin (Hs-MCU, Nc-MCU, Af-MCU, ^HsMTS^Af-MCU, ^HsMTS^Nc-MCU, Hs-MICU1, and Hs-MICU1^mEF^ overexpressing HeLa or HEK-293 cells) at 37°C and 5% CO_2_. The lentiviral vector pLX304 was obtained from the Broad Institute’s RNAi Consortium and used for expressing V5-tagged cDNAs. All chemicals were purchased from Sigma-Aldrich, unless specified.

### Isolation of Crude HeLa Mitochondria

Cell lysates and crude mitochondria were prepared from cultured HeLa cells as previously described (Wettmarshausen and Perocchi, 2017) and analyzed by alkaline carbonate extraction.

### Measurements of Mitochondrial Calcium Uptake in HeLa Cells

Mitochondrial Ca^2+^ uptake was measured by mt-AEQ-based measurements of Ca^2+^-dependent light kinetics upon 100 μM histamine stimulation as previously described (Arduino et al., 2017).

### Yeast Strains

Yeast strains expressing mt-AEQ or cyt-AEQ were generated by transforming the yeast wild-type YPH499 strain and selecting transformants in glucose medium lacking uracil (Sikorski and Hieter, 1989). Yeast strains expressing different combinations of Hs-EMRE, Hs-MCU, Hs-MICU1 and their mutants were generated by simultaneously transforming the YPH499 strain with the respective plasmids and by selecting transformants on glucose medium lacking uracil (AEQ), histidine (Hs-MCU), leucine (Hs-EMRE), and tryptophan (Hs-MICU1) as selection markers.

### Measurements of Calcium Kinetics in Yeast

*In vivo* analyses of cytosolic and mitochondrial Ca^2+^ kinetics in yeast cells were performed by mt-AEQ and cyt-AEQ-based measurements of Ca^2+^-dependent aequorin light response upon CaCl_2_ (1 mM) and glucose addition (100 mM). Ca^2+^ uptake by mitochondria isolated from yeast strains expressing Hs-MICU1 was measured with Calcium Green-5N upon CaCl_2_ injection (100 μM final concentration).

### Yeast Growth Measurement

Yeast cultures were grown for 48-72 hours in a selective lactate medium (S-LAC) under different environmental stresses and their doubling time was calculated.

### MTT Assay

A colorimetric 3-(4,5- dimethylthiazol- 2- yl)-2,5- diphenyltetrazolium bromide (MTT) metabolic activity assay was used to determine cell viability.

### Quantification and Statistical Analysis

Data are represented as mean ± SEM and the statistical analysis of each experiment is described in the figure legends. Normal distribution was tested by Shapiro-Wilk normality test. Differences between two datasets were evaluated by two-tailed unpaired Student’s t test. Statistical tests between multiple datasets and conditions were carried out using one-way analysis of variance (ANOVA) followed by Tukey’s or Dunnett’s Multiple Comparison. Statistical analyses were performed using GraphPad Prism.

## ACKNOWLEDGEMENTS

We thank Dr. Daniela M. Arduino for critical reading of the manuscript. We acknowledge support from the German Research Foundation (DFG) under the Emmy Noether Programme (PE 2053/1-1 to F.P. and J.W.); the Munich Center for Systems Neurology (SyNergy EXC 1010 to F.P.); the Juniorverbund in der Systemmedizin ‘mitOmics’ (FKZ 01ZX1405B to V.G. and A.L.); The Bert L & N Kuggie Vallee Foundation (to F.P.); the DFG (MO1944/1-2 to D.M.); The Spanish Ministry of Economy, Industry, and Competitiveness (MEIC; BFU2015-67107) and from the European Union’s Horizon 2020 research and innovation programme under the grant agreement ERC-2016-724173 (to T.G. and A.P.).

## SUPPLEMENTAL INFORMATION

Supplemental Information includes Supplemental Experimental Procedures.

## AUTHOR CONTRIBUTIONS

Conceptualization, F.P.; Methodology, V.G., J.W., D.M., T.G.; Formal Analysis, V.G., J.W., A.A.P., A.L., Y.C.; Resources, F.P., D.M.; Writing-Original Draft, F.P., V.G., J.W.; Visualization, F.P., V.G., J.W.; Supervision, F.P.; Funding Acquisition, F.P., D.M., and T.G.

## SUPPLEMENTAL EXPERIMENTAL PROCEDURES

### Phylogenetic Profiling of MICU1 and MCU

Homologs of human MCU and MICU1 across 247 eukaryotes were retrieved from ProtPhylo (www.protphylo.org) (Cheng and Perocchi, 2015) using OrthoMCL with more than 0% match length and inflation index of 1.1 for orthology assignment. The percentage of amino acids match length was determined based on blastp. The phylogenetic tree of 247 eukaryotes was reconstructed using the phylogenetic tree generator (https://phylot.biobyte.de/) and visualized using iTOL (https://itol.embl.de/). The mitochondrial-targeting sequence (MTS) probability was determined with MitoProt (https://ihg.gsf.de/ihg/mitoprot.html).

### Protein Domains

Protein sequences of *Homo sapiens* MCU (Hs-MCU, NP_612366.1) *Neurospora crassa* MCU (Nc-MCU, XP_959658.1), and *Aspergillus fumigatus* MCU (Af-MCU, XP_751795.1) were analyzed to predict MTS, DUF607 motif (Finn et al., 2016), coiled coil domains (CCD) (https://embnet.vital-it.ch/software/COILS_form.html), and transmembrane domains (TM) (TMHMM 2.0). Clustal Omega was used for proteins alignment and sequence similarities above 80% were color-coded with the Sequence Manipulation Suite.

### Plasmids and Reagents

The lentiviral vector pLX304 was obtained from the Broad Institute’s RNAi Consortium and used for expressing V5-tagged cDNAs. Full-length, human wild-type EMRE (Hs-EMRE), MCU (Hs-MCU), MICU1 (Hs-MICU1) and their mutants (Hs-MCU_D261A_, Hs-MCU_E264A_, and Hs-MICU1_mEF_) cDNAs without a stop codon were obtained from Addgene. Hs-MICU1_mEF_ contains two point mutations in the first (D231A, E242K) and the second EF-hand domains (D421A, E432K) as described in (Perocchi et al., 2010). Af-MCU and Nc-MCU with (^HsMTS^Af-MCU and ^HsMTS^Nc-MCU) and without the N-terminal MTS of Hs-MCU (HsMTS, aminoacids 1-56) and without a stop codon were codon optimized for human expression and synthesized *de novo* in the PuC57 vector (GenScript). Cytosolic aequorin (cyt-AEQ) was kindly provided by Teresa Alonso and a mitochondria-targeted aequorin (mt-AEQ) was generated as previously described in (Arduino et al., 2017). Hs-EMRE, Hs-MCU, Hs-MICU1 and their mutants cDNAs were amplified by PCR using the following primers: fw-MCU, 5’-CCC TCT AGA ATG GCG GCC GCC GCA GGT AG-3’; rv-MCU, 5’-GGG CTC GAG TTA ATC TTT TTC ACC AAT TTG TCG-3’; fw-MICU1, 5’-CCC GGA TCC ATG TTT CGT CTG AAC TCA CTT TC-3’; rv-MICU1, 5’-GGG CTC GAG TTA CTG TTT GGG TAA AGC GAA G-3’; fw-EMRE, 5’-CCC GGA TCC ATG GCG TCC GGA GCG GCT CGC-3’; rv-EMRE, 5’-GGG CTC GAG TTA GTC ATC ATC ATC ATC ATC CTC-3’). PCR products were cloned into the yeast expression plasmids p423GPD (Hs-MCU, Hs-MCU_D261A_, Hs-MCU_E264A_), p414GPD (Hs-MICU1, Hs-MICU1_mEF_) and p425GPD (Hs-EMRE) (Mumberg et al., 1995). For expression in mammalian cells, cDNAs were first cloned into the pDONR221 Gateway vector (Thermo Fisher Scientific, 1253607) and subsequently transferred into the pLX304 vector according to manufacturer’s instructions (Life Technologies). All chemicals were purchased from Sigma-Aldrich, unless specified.

### Cell Lines

All mammalian cells were grown in high-glucose Dulbecco’s modified Eagle’s medium (DMEM) supplemented with 10% FBS at 37°C and 5% CO_2._ HeLa cells stably expressing mt-AEQ were generated as previously described (Arduino et al., 2017) and maintained in 100 μg/ml geneticin (Thermo Fisher Scientific, 10131027). Mt-AEQ HeLa cells stably overexpressing either a shRNA targeting Hs-MCU (shMCU; Sigma Aldrich, TRCN0000133861) or an empty vector (pLKO; Addgene, 8453) were generated as previously described (Baughman et al., 2011) and maintained in 2 μg/mL puromycin (Life Technologies, A11138) and 100 μg/ml geneticin. shMCU, mtAEQ HeLa cells stably overexpressing Hs-MCU, Nc-MCU, Af-MCU, ^HsMTS^Af-MCU and ^HsMTS^Nc-MCU from the pLX304 lentiviral vector were generated by transduction. Lentivirus production and infection were performed according to guidelines from the Broad RNAi Consortium and infected cell lines were selected with 10 μg/mL blasticidin (Life Technologies, R21001), 2 μg/mL puromycin, and 100 μg/ml geneticin. MICU1-knockout HEK-293 cells were kindly provided by Vamsi Mootha. MICU1-knockout HEK-293 cells stably overexpressing either wild-type or mutant Hs-MICU1 (Hs-MICU1_mEF_) from the pLX304 vector were generated by transduction and selected with 10 μg/mL blasticidin.

### Isolation and Analysis of Crude HeLa Mitochondria

Cell lysates and crude mitochondria were prepared from cultured HeLa cells as previously described (Wettmarshausen and Perocchi, 2017) and immunoblotted with the following antibodies: anti-MCU (Sigma Aldrich, HPA01648), anti-V5 (Life Technologies, R96025), and anti-ATP5A (Abcam, MS507).

Alkaline carbonate extraction from crude mitochondria was performed as described previously (Baughman et al., 2011). Mitochondrial soluble and membrane fractions were immunoblotted with anti-V5, anti-TIM23 (BD Bioscience, 611222), and anti-HSP60 (R&D System, MAB1800) antibodies.

### Measurements of Mitochondrial Calcium Uptake in HeLa Cells

Mitochondrial Ca^2+^ uptake was measured in mt-AEQ HeLa cells as previously described (Arduino et al., 2017). Briefly, HeLa cells stably expressing mt-AEQ were seeded in white 96-well plates at 25,000 cells/well in growth medium. After 24 hours, mt-AEQ was reconstituted with 2 μM native coelenterazine (Abcam, ab145165) for 2 hours at 37°C. Mt-AEQ-based measurements of Ca^2+^-dependent light kinetics were performed upon 100 μM histamine stimulation. Light emission was measured in a luminescence counter (MicroBeta2 LumiJET Microplate Counter, PerkinElmer) at 469 nm every 0.1 s. At the end of each experiment, cells were lysed with a solution containing 0.1 mM digitonin and 10 mM CaCl_2_ to release all the residual aequorin counts. Quantification of mt-Ca^2+^ concentrations was performed using the algorithm reported in (Bonora et al., 2013).

### Yeast Strains

Yeast strains expressing mt-AEQ or cyt-AEQ were generated by transforming the yeast wild-type YPH499 strain and selecting transformants in glucose medium lacking uracil (Sikorski and Hieter, 1989). Yeast strains expressing different combinations of Hs-EMRE, Hs-MCU, Hs-MICU1 and their mutants were generated by simultaneously transforming the YPH499 strain with the respective plasmids and by selecting transformants on glucose medium lacking uracil (AEQ), histidine (Hs-MCU), leucine (Hs-EMRE), and tryptophan (Hs-MICU1) as selection markers. To test expression and mitochondrial localization of heterologous proteins, yeast were grown at 30°C in a selective lactate medium (S-LAC) containing 8.5 g/L yeast nitrogen base, 25 g/L ammonium sulfate, 2% (v/v) lactic acid, 0.1 % glucose, supplemented with their respective selection markers at pH 5.5/KOH. At an OD ~0.8 cells were harvested at 1000 g for 5 min at room temperature. The cell pellet was re-suspended in SHK buffer (0.6 M sorbitol, 20 mM HEPES/KOH pH 7.2, 80 mM KCl, and 1 mM PMSF) and vortexed with glass beads (425-600 μm diameter) 5 times for 30 s with cooling down in between. This mix was centrifuged at 1000 g for 5 min at 4°C and the supernatant was further centrifuged at 20,000 g for 10 min at 4°C. The resulting supernatant (cytosolic fraction) was precipitated with trichloroacetic acid. Both cytosolic and mitochondrial (pellet) fractions were directly resuspended in Laemmli buffer and separated under reducing conditions on 12 or 14% SDS-PAGE gels. Immunoblotting was performed according to standard procedures using the following antibodies: anti-MCU (Sigma-Aldrich, HPA016480); anti-EMRE (Santa Cruz Biotechnology, sc-86337); anti-MICU1 (Sigma Aldrich, HPA037480); anti-YME1 (Thermofisher/Novex, 459250); AEQ (Merck/Millipore, MAB4405).

### Measurements of Calcium Kinetics in Yeast

*In vivo* analyses of cytosolic and mitochondrial Ca^2+^ kinetics in yeast cells were performed as described by (Groppi et al., 2011) with some modifications. Exponentially growing cells (OD 0.8, ~24×10^6^ cells/mL) in S-LAC at 30°C were harvested by centrifugation at 3,500 g for 5 min at room temperature. The yeast cell pellet was washed three times with RT milliQ water and then resuspended in a nutrient-free buffer (NFB; 100 mM Tris, pH 6.5) at a density of 1×10^8^ cells/mL. Cells were incubated for 1.5 h at room temperature, collected by centrifugation at 3,500 rpm for 5 min and concentrated in the same buffer at a density of 25×10^8^ cells/mL. The photoprotein aequorin was then reconstituted with 50 μM native coelenterazine for 30 min at RT in the dark. Excess of coelenterazine was removed by three washes with NFB and the cell pellet was then resuspended at a final density of 5×10^8^ cells/mL. About 0.5×10^8^ cells/well were placed in a white 96-well plate and Ca^2+^-dependent aequorin light response was measured upon addition of a spiking solution containing CaCl_2_ and glucose to a final concentration of 1 mM and 100 mM, respectively, at 0.5 s intervals in a MicroBeta2 LumiJET Microplate Counter. At the end of each experiment, a lysis solution containing 5 mM digitonin, 450 mM EGTA, 100 mM Tris (pH 6.5/KOH) was added at a ratio pf 1:5 for 5 min at 37°C and light response was measured upon addition of CaCl_2_ to a final concentration of 100 mM to release all the residual aequorin counts. Quantification of mt-Ca^2+^ concentrations was performed using the algorithm reported in (Bonora et al., 2013).

Measurements of Ca^2+^ uptake by mitochondria isolated from yeast strains expressing Hs-MICU1 were performed as it follows: crude mitochondria were isolated as described previously (Arduino et al., 2017) and 150 μg/well were placed in a black 96-well plate and incubated in a buffer containing 0.6 M sorbitol, 20 mM HEPES, 2 mM MgCl_2_, 10 mM KH_2_PO_4_, 3 mM glutamate, 3 mM malate, 3 mM succinate, 50 μM EDTA, and 0.1 μM Calcium Green-5N (Life technologies, C3737) for 3 min. Ca^2+^ uptake was initiated by CaCl_2_ injection (100 μM final concentration) and extramitochondrial free Ca^2+^ was monitored every 2 s at room temperature using a CLARIOstar microplate reader (BMG Labtech) at 506 and 531 nm excitation and emission, respectively. The MCU inhibitor Ru360 (10 μM, final concentration) was used as a positive control.

### Yeast Growth Measurement

Overnight yeast cultures grown at 30°C in S-LAC were diluted to an OD of 0.1 (3×10^6^ cells/mL) and then 0.3×10^6^ cells/well were placed in a black, gas-permeable Lumox 96-well plate. The yeast suspension was measured at an absorbance of 600 nm and an interval of 340 s in the CLARIOstar microplate reader (BMG Labtech) for 48-72 hours with shaking at 30°C, 37°C, or in presence of sterile solutions of sodium chloride (NaCl, 0.1-1 M), copper (II) chloride (CuCl_2_, 10-30 mM), iron (II) chloride (FeCl_2_, 10-40 mM), manganese (II) chloride (MnCl_2_, 1-5 mM), strontium (II) chloride (SrCl_2_, 10-50 mM), zinc (II) chloride (ZnCl_2_, 10-50 mM), or antifungal drugs (miconazole, 10-100 ng/ml; amoidarone, 5-20 μM). The average time taken by the yeast culture to double in the log-growth phase (doubling time) was calculated using the following equation:

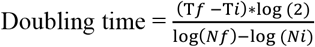

where T is the time between the log-growth phase from T*i* to T*f* and N is the number of cells measured as an optical density at an absorbance of 600 nm at the time point T*i* (N*i*) and T*f* (N*f*).

### MTT Assay

A colorimetric 3-(4,5-dimethylthiazol-2-yl)-2,5-diphenyltetrazolium bromide (MTT) metabolic activity assay was used to determine cell viability. HEK-293 cells were seeded at 50,000 cells/well in 1 mL of DMEM with high glucose and 10% FBS in a transparent 24-well plate at 37 °C and 5% CO_2_. After 24 hours, FeCl_2_ (0.1-1 mM) or MnCl_2_ (5-50 μM) were added to the well and the cells were incubated further for 48 h. Afterwards, 500 μL of medium was removed from the well, 50 μl of MTT solution (Sigma Aldrich, M5655; 5 mg/ml in PBS) was added, and cells were incubated for 3 h at 37 °C. Finally, cells were lysed with 500 μl of solubilization solution (1% SDS and 0.1 M HCl in isopropanol) for 15 min at 37 °C and absorbance was monitored in a CLARIOstar microplate reader (BMG Labtech) at 570 nm.

### Data Analysis

All data are represented as mean ± SEM and the statistical details of experiments can be found in the figure legends. Normal distribution was tested by Shapiro-Wilk normality test. Differences between two datasets were evaluated by two-tailed unpaired Student’s t test. Statistical tests between multiple datasets and conditions were carried out using one-way analysis of variance (ANOVA) followed by Tukey’s or Dunnett’s Multiple Comparison. Statistical analyses were performed using GraphPad Prism.

